# Mechanisms of a locally adaptive shift in allocation among growth, reproduction, and herbivore resistance in *Mimulus guttatus*

**DOI:** 10.1101/400523

**Authors:** David B. Lowry, Damian Popovic, Darlene J. Brennan, Liza M. Holeski

**Author notes:** Corresponding author: David B. Lowry.

## Abstract

Environmental gradients can drive adaptive evolutionary shifts in plant resource allocation among growth, reproduction, and herbivore resistance. However, few studies have attempted to connect these adaptations to underlying physiological and genetic mechanisms. Here, we evaluate potential mechanisms responsible for a coordinated locally adaptive shift between growth, reproduction, and herbivore defense in the yellow monkeyflower, *Mimulus guttatus.* Through manipulative laboratory experiments we found that gibberellin (GA) growth hormones may play a role in the developmental divergence between perennial and annual ecotypes of *M. guttatus*. Further, we detected an interaction between a locally adaptive chromosomal inversion, *DIV1*, and GA addition. This finding is consistent with the inversion contributing to the evolutionary divergence between inland annual and coastal perennial ecotypes by reducing GA biosynthesis/activity in perennials. Finally, we found evidence that the *DIV1* inversion is partially responsible for a coordinated shift in the divergence of growth, reproduction, and herbivore resistance traits between coastal perennial and inland annual *M. guttatus*. The inversion has already been established to have a substantial impact on the life-history shift between long-term growth and rapid reproduction. Here, we demonstrate that the *DIV1* inversion also has sizable impacts on both the total abundance and composition of phytochemical compounds involved in herbivore resistance.

## INTRODUCTION

One of the fundamental tenets of evolutionary biology is that adaptation of organisms to specific environmental conditions inevitability results in a fitness trade-off. Trade-offs often manifest in the form of a cost, such that organisms that become adapted to one set of environmental conditions will be at a disadvantage in alternative environments (Futuyma and Moreno 1988; Whitlock 1996). The idea of trade-offs involved in adaptation and ecological specialization has been borne out in a wide range of evolutionary scenarios, including predator-prey relationships and host-races formation in insect herbivores (Futuyma and Moreno 1988; Kawecki 1998; Svanback and Eklov 2003; Forister et al. 2012). A common source of ecological specialization and consequent trade-offs is the process of local adaptation across environmental gradients (Kawecki and Ebert 2004; Hereford 2009).

Local adaption across environmental gradients can lead to shifts in the allocation of resources to long-term growth (survival) and reproduction (fecundity; Clausen and Hiesey 1958; Lowry 2012; Friedman and Rubin 2015). Those shifts in life-history strategy along environmental gradients can also have major impacts on allocation to herbivore defense (Hahn and Maron 2016). However, there appears to be key differences in how resources are allocated across environmental gradient for interspecific and intraspecific comparisons. Interspecific variation in plant species typically fits well with the resource allocation hypothesis (Coley et al. 1985), where low resource environments tend to be composed of slower growing better defended species while high resource environments promote faster growing poorly defended plants (Endara and Coley 2011). In contrast, intraspecific variation along environmental gradients is far less consistent and more often than not contradicts the predictions of the resource allocation hypothesis (Hahn and Maron 2016). One common pattern for intraspecific plant variation is a positive relationship between the length of the growing season along an environmental gradient and the level of herbivore resistance (Hahn and Maron 2016; Kooyers et al. 2017). This pattern could be driven by plants having more time to allocate resources to leaf production and defense, greater herbivore pressure in habitats with longer growing seasons, and/or a greater apparency of plants with longer growing seasons (Feeny 1976; Mason and Donovan 2015; Hahn and Maron 2016; Kooyers et al. 2017). Another important factor to consider is allocation to reproduction (fecundity), which frequently trades off with constitutive and/or induced herbivore resistance (Agren and Schemske 1993; Heil and Baldwin 2002; Strauss et al. 2002; Stowe and Marquis 2011; Cipollini et al. 2014).

Achieving an evolutionary optimum in how resources are allocated to growth, reproduction, and defense will depend on the nature of all environmental challenges faced by each local population (Rhoades 1979; Rausher 1996; Hamilton et al. 2001; Strauss et al. 2002; Stamp 2003; Karban 2011; Cipollini et al. 2014; Jensen et al. 2015; Smilanich et al. 2016). Despite the development of multiple ecological and evolutionary hypotheses that postulate a relationship between growth, reproduction, and resistance to herbivores (Feeny 1976; Coley et al. 1985; Rhoades 1979; Herms and Mattson 1992; Strauss et al. 2002; Stamp 2003, Fine et al. 2006; Agrawal et al. 2010; Cipollini et al. 2014; Hahn and Maron 2016), these hypotheses do not make any predictions about the underlying molecular mechanisms that mediate these relationships. The genetic mechanisms responsible for trade-offs among growth, reproduction, and resistance are just beginning to be elucidated in model systems (Lorenzo et al. 2004; Yang et al. 2012; Kerwin et al. 2015; Campos et al. 2016; Havko et al. 2016; Major et al. 2017; Howe et al. 2018; Rasmann et al. 2018), but have yet to be evaluated in the evolutionary context of local adaptation.

Recent studies have shown that changes in the allocation of resources to growth versus resistance are made through a set of interacting gene networks (Karzan and Manners 2012; Huot et al. 2014; Campos et al. 2016; Havko et al. 2016). Jasmonates (JA) are key regulatory hormones involved in the response of plants to herbivore attack (Zhang and Turner 2008; Havko et al. 2016). While JA production increases herbivore defenses, it also inhibits plant growth through interactions with other gene networks (Zhang and Turner 2008; Yan et al. 2007; Karzan and Manners 2012; Yang et al. 2012). For example, the critical cross-talk between downstream genes in the JA pathway (JAZ genes) and Gibberellin (GA) pathway (DELLA genes) are thought to play a pivotal role in mediating resource allocation (Yang et al. 2012; Hou et al. 2013; Havko et al. 2016).

Here, we focus on understanding the physiological and genetic mechanisms underlying shifts in allocation to growth, reproduction, and defense for local adapted populations of the yellow monkeyflower *Mimulus guttatus*. The availability of soil water is a key driver of local adaptation in the *M. guttatus* species complex (Hall and Willis 2006; Lowry et al. 2008; Ferris et al. 2017). The coastal habitats of California and Oregon have many wet seeps and streams that are maintained year-round as a result of persistent summer oceanic fog and cool temperatures (Hall and Willis 2006; Lowry et al. 2008; Lowry and Willis 2010). All coastal populations of *M. guttatus* that reside in those habitats have a late-flowering life-history strategy. These coastal perennial populations thus, make a long-term investment in growth over reproduction (Hall and Willis 2006; Lowry et al. 2008; Hall et al. 2010; Baker and Diggle 2011; Baker et al. 2012). That investment in growth manifests through the production of many vegetative lateral stolons, adventitious roots, and leaves in coastal perennial plants. In contrast, the vast majority of nearby inland populations of *M. guttatus* in the coastal mountain ranges reside in habitats that dry out completely during summer months. These inland populations have evolved a rapid growth, drought escape annual life-history strategy. Instead of investing in vegetative lateral stolons, the axillary branches of inland plants are mostly upcurved and typically produce flowers quickly (Lowry et al. 2008; Lowry and Willis 2010; Friedman et al. 2015). Further, inland plants invest less into the production of leaves before flowering than coastal perennials (Friedman et al. 2015). It should be noted that a smaller number of inland populations in the coastal mountain ranges do reside in rivers and perennials seeps and have a perennial life-history. Farther inland, perennial populations are more common, especially in high elevation streams and hot springs of the Sierra and Cascade Mountains (Oneal et al. 2014). The inland perennial plants are similar to coastal perennials in overall morphology, especially in that they are more prostrate in growth habit, with the production many stoloniferous branches. However, the coastal perennials differ from inland perennial by being more robust in their growth form and having evolved tolerance to oceanic salt spray (Lowry et al. 2009).

While perennial populations invest more into vegetative growth than reproductive growth, they also invest more heavily in defending their vegetative tissues. Perennial populations of *M. guttatus* have higher levels of both constitutive and induced defensive phenylpropanoid glycoside (PPG) compounds than the annual populations when grown in a common environment (Holeski et al. 2013). PPG levels have a negative relationship with the performance of multiple generalist herbivores in *M. guttatus* (Rotter et al. 2018). This pattern of highly defended plants in wetter habitats with long growing seasons versus poorly defended plants in dry habitats with short growing seasons is consistent with an optimal defense strategy: Greater allocation of resources to herbivore resistance is favored in habitats with a long growing season by both a greater abundance of herbivores and a lower physiological cost of producing defensive compounds due to high soil water availability (Kooyers et al. 2017).

Two major QTLs (*DIV1* and *DIV2*) and many minor QTLs control key traits involved in local adaptation to perennial and annual habitats within the *M. guttatus* species complex (Hall et al. 2006; Lowry and Willis 2010; Hall et al. 2010; Friedman and Willis 2013; Friedman et al. 2015). *DIV1* has the largest effect on the most traits and has thus been more extensively studied than *DIV2*. *DIV1* is a large paracentric chromosomal inversion that plays a pivotal role in the annual versus perennial life-history divergence described above (Lowry and Willis 2010). The inversion is at minimum 6.3 Mbp in length along linkage group 8 (LG8) and contains at least 785 annotated genes. In hybrids, *DIV1* has a major effect on growth rate including the adaptive flowering time phenotype, explaining 21% to 48% of the divergence between inland annual and coastal perennial parents (Lowry and Willis 2010). In addition to flowering time, the *DIV1* inversion has major effects on multiple traits involved in the evolutionary shift from more allocation of resources to long-term growth versus reproduction (Lowry and Willis 2010). Plants with the perennial (PE) orientation of the inversion have a more prostrate growth habitat, produce more lateral stolons and adventitious roots, have thicker shoots, and larger flowers than plants with the annual (AN) orientation of the inversion (Lowry and Willis 2010; Friedman et al. 2015). A recent allele frequency outlier analysis of coastal perennial and inland annual populations identified candidate genes in the gibberellin pathway that may underlie a pleiotropic shift in allocation between growth and reproduction (Gould et al. 2017). That study also found evidence of recent selection on a key GA biosynthetic gene, *GA20-oxidase2* (*GA20ox2*), within the *DIV1* inversion and for the *Gibberellic Acid Insensitive* (*GAI*) in the vicinity of *DIV2.*

Given the recent outlier analysis identifying divergence among genes in the GA pathway and the GA pathways interaction with the JA herbivore resistance pathway, we evaluate the role of GA in the divergence of growth morphology and herbivore resistance between perennial and annual ecotypes of *M. guttatus.* We hypothesized that perennial accessions would display a greater morphological response to GA addition than annual accession, as their prostrate morphology was consistent with lower GA biosynthesis and/or downstream signaling activity. Further, we hypothesized that the adaptive *DIV1* inversion contributes to the lower GA biosynthesis/activity in coastal perennial plants. To test this hypothesis, we utilized near-isogenic lines (NILs) for the *DIV1* locus and evaluated whether there was interaction between the *DIV1* inversion and GA addition. If the GA biosynthesis/activity is down regulated by the perennial orientation of the inversion, then we predict that the NILs containing the perennial inversion orientation will respond more to the GA addition than the NILs containing the annual orientation. Finally, we tested the hypothesis that the *DIV1* inversion is partially responsible for the evolution of locally adaptive trade-offs in allocation between reproduction and defense that has been broadly observed for populations of *M. guttatus* that vary in growing season length (Lowry et al. 2008; Holeski et al. 2013; Kooyers et al. 2017). To test this hypothesis, we compared the concentrations of PPGs between NILs containing coastal perennial and inland annual orientations of the *DIV1* inversion.

## METHODS

### Plant material

For comparisons among ecotypes, we utilized single family population accessions derived from coastal perennial, inland annual, and inland perennial populations of *M. guttatus* (Fig. 1). The locations from where population accessions were collected are listed in Table S1. Previous population structure analyses found that coastal perennial populations of *M. guttatus* are more closely related to each other than they are to the inland annual populations (Lowry et al. 2008; Twyford and Friedman 2015). Thus, the coastal populations collectively constitute a distinct locally adapted ecotype (Lowry 2012). In contrast, population structure between inland annuals and inland perennial populations is generally low (Twyford and Friedman 2015). However, particular regions of the genome, including an adaptive chromosomal inversion (*DIV1*, discussed below) are more differentiated between inland annuals and inland perennials (Oneal et al. 2014; Twyford and Friedman 2015). We therefore consider inland annuals and inland perennials as different ecotypes as well.

**Figure 1.**
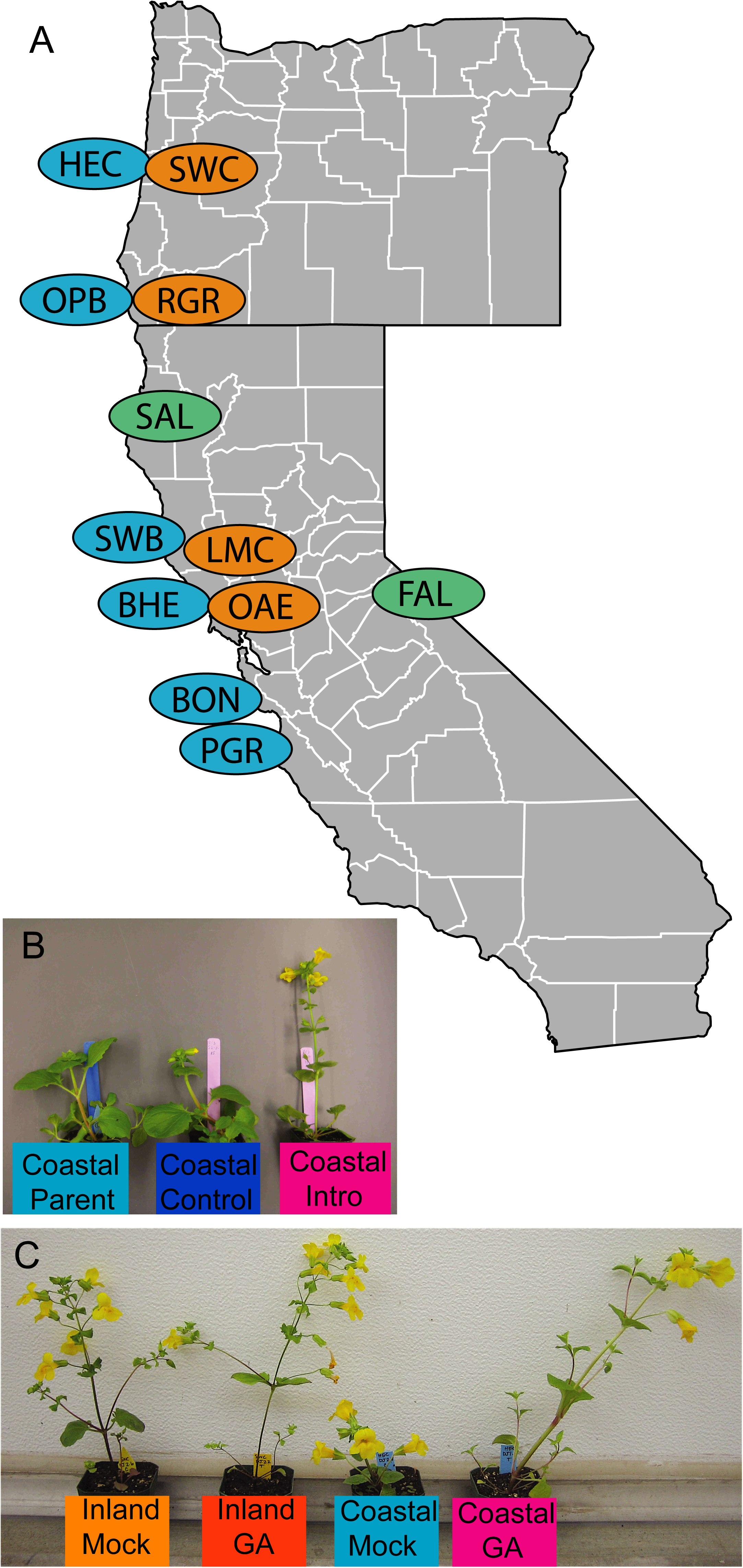
A) Map of coastal perennial (blue), inland annual (yellow), and inland perennial (green) populations from where accessions for experiment were derived. B) Effects of the introgression of the inland *DIV1* inversion orientation into the coastal perennial genetic background. Left: Coastal perennial parental inbred line. Center: Control Near Isogenic Line (NIL) containing the coastal orientation of the inversion. Right: Introgression NIL containing the inland orientation of the inversion. All three plants germinated on the same day. C) Differential responses of inland annual and coastal perennial accessions to gibberellin treatment and water control (mock).

To understand the phenotypic effects of the *DIV1* inversion, Lowry and Willis (2010) previously created near-isogenic lines (NILs) for the inversion. The NILs are the product of crosses between inbred lines from the coastal perennial SWB population and the nearby inland annual LMC population. F1 hybrids were recurrently backcrossed to both of their respective parents for four generations. Heterozygous fourth generation backcrosses were then self-fertilized to produce two types of NILs: 1) Individuals that were homozygous for the introgressed allele of *DIV1* (Introgression-NILs) and 2) Individuals that were homozygous for the *DIV1* allele of the genetic background (henceforth referred to as Control-NILs). Comparisons between Introgression-NILs and Control-NILs are ideal for testing inversion function because their genetic backgrounds are nearly identical, but they are homozygous for opposite *DIV1* alleles.

### The effects of GA application on plant growth among ecotypes

To evaluate whether perennial and annual plants differ in their response to GA addition, we conducted a greenhouse experiment with accessions derived from five coastal perennial, four inland annual, and two inland perennial populations (Table S1). Seeds were sown in Suremix soil (Michigan Grower Products Inc., Galsburg, MI) and stratified at 4° C for two weeks. After stratification, pots were moved to the Michigan State University greenhouses. Temperature was set in the greenhouse room to 22° C days/18° C nights. Plants were grown in 16-hour days and 8-hour nights, where supplemental lighting was used during the full day period. Seedlings were transplanted to individual 3.5-inch square pots filled with Suremix soil. Transplanted seedlings were randomized across the experiment and randomly assigned to a GA treatment group or a mock control group. After transplantation, plants were sprayed five times each, every other day, with 100mM GA3 (GA treatment) or DI water (mock). This daily spray volume amounts to ∼3.5 mL.

To standardize the developmental time point at which plant traits were quantified, we measured the following traits on the day of first flowering: Total number of nodes on the primary shoot, lengths and widths of the first three internodes, length and width of the corolla of the first open flower, plant height, the total number of adventitious roots at the first node of all branches, total number of stoloniferous nodes sprouting adventitious roots, total number of aerial branches, total number of stolons, length of the longest aerial branch, length of the longest stolon, and the length and width of the longest leaf at the second node. Ten days after first flower, we quantified the same traits as at first flower, with the following exceptions: Length and width of corollas were not quantified, but we did count the total number of flowers.

Results were analyzed with JMP 14.0.0 (SAS Institute, Cary, NC). To test the hypothesis that the perennials respond morphologically more to the addition of GA, we conducted principal components analyses using all of the measured traits. Since many of these traits were highly correlated (Table S1), we reduced the dimensionality of the data with a principal components (PC) analysis. We conducted the PC analysis on correlations in JMP. We used a Scree Test (Cattell 1966) to determine how many principal components to retain. To understand the effects of accession, ecotype, GA treatment, and the interactions of these effects on each PC, we fit the default standard least squares ANOVA model with the Fit Model platform in JMP. Each of the PCs were modeled as response variables to the following fixed-effect factors and interactions: accession (nested within ecotype), ecotype (coastal perennial, inland annual, inland perennial), treatment (GA vs. Mock), accession x treatment, and ecotype x treatment. A significant ecotype x treatment interaction would indicate that the ecotypes vary in their response to GA. Further, if perennials responded more to the GA treatment it would support the hypothesis that the perennials have less GA biosynthesis/activity than the annuals.

### Interactions of GA application with the adaptive DIV1 inversion

We grew coastal (SWB-S1) and inland (LMC-L1) parental inbred lines along with the NILs derived from those lines in a fully randomized design in the Michigan State University greenhouses, with 16-hours of supplemental lighting. We focused on the effect of the inversion in the coastal perennial genetic background, as a previous study had shown that the effect of the inversion had the greatest effect in the perennial genetic background (Friedman 2014). Following transplantation, seedlings were sprayed with GA or a mock water treatment every other day and traits were quantified in the same way as for the comparing population accessions.

To establish how trait variation of the coastal and inland parental lines was influenced by the GA treatment, we conducted a PC analysis of all plants in the experiment in JMP 14.0.0. As for the analysis with multiple population accessions (above), we conducted principal components analyses using all traits and used a Scree Test (Cattell 1966) to determine how many principal components to retain. Models were fit for each PC to test the effect of the following fixed-effect factors: line, treatment, and the line x treatment interaction. We fit models independently for the parental lines and the NILs. The analysis of the parental lines was conducted to confirm that the responses to GA were similar as for the population survey (above). The analysis of the NILs allowed us to test whether there was a significant interaction of the inversion with the GA treatment (i.e. line x treatment interaction). Line x treatment least square means were compared with Tukey HSD post hoc analyses for both parental lines and NILs. A significant line x treatment interaction for the NILs, with a greater response for the introgression NILs, would support the hypothesis that the perennial orientation of the *DIV1* inversion reduces GA biosynthesis/activity.

### Effects of the DIV1 inversion on resistance compound concentrations

To evaluate the effects of the *DIV1* inversion on the production of herbivore resistance compounds, we conducted an experiment using the inversion NILs. Seeds were stratified, germinated, and transplanted following the same protocols as in the previous two experiments. In contrast to the GA NIL experiment, we grew outbred NILs which were created by intercrossing independently derived NILs because we did not have any prior data on the impacts of inbreeding on PPG production. As for the GA experiment, we focused our study on the effect of the inversion in the perennial genetic background. Here, we used the outbred NILs made by intercrossing SWB-S2 and SWB-S3 derived coastal genetic background NILs. We used intercrosses between LMC-L2 andLMC-L3 for the inland parents and between SWB-S2 and SWB-S3 for the coastal perennial parent comparisons. See Lowry & Willis (2010) for full description of outbred NIL generation.

To ensure that enough leaf tissue was available for analyses, outbred NILs and outbred parents were allowed to flower prior to the collections for PPG quantification. Collected leaf tissue was lyophilized for two days and then shipped to Northern Arizona University for analyses. We ground the leaf tissue using a 1600 MiniG (Spex, Metuchen, New Jersey). Extractions were conducted in methanol, as described in Holeski et al. 2013, 2014. We quantified PPGs using high performance liquid chromatography (HPLC), via an Agilent 1260 HPLC (Agilent Technologies, Santa Clara, California) with a diode array detector and Poroshell 120 EC-C18 analytical column (4.6 x 250mm, 2.7µm particle size) maintained at 30°C. HPLC run conditions, were conducted as described in Kooyers et al. (2017). We calculated concentrations of PPGs as verbascoside equivalents, using a standard verbascoside solution (Santa Cruz Biotechnology, Dallas, Texas), as described in Holeski et al. (2013, 2014). We compared the concentrations of total PPGs and individual PPGs with one-way ANOVAs fit in JMP 12.2.0. Post-hoc Tukey HSD Tests were used to compare means of parental and NIL classes.

## RESULTS

### The effects of GA application on plant growth among ecotypes

Consistent with previous studies (Hall et al. 2006; Lowry et al. 2008; Lowry and Willis 2010; Oneal et al. 2014), there were large differences in morphology between coastal perennial, inland annual, and inland perennial ecotypes. The perennials, especially the coastal ones, were generally larger overall across a suite of traits. The perennials also all had a prostrate growth habit with many lateral stolon branches that produced adventitious roots. In contrast, the inland annuals had thinner shoots, smaller flowers, and primarily produced upcurved aerial branches lacking adventitious roots. We reduced the dimensionality of the data using a principal components analysis, retaining the first 4 PCs based on a Scree Test (Figure S1; Cattell 1966). The first four PCs collectively explained 70.3% of the phenotypic variation. Most traits (32 out of 37) heavily loaded (Loadings > 0.40) onto the first PC (Eigenvalue = 14.698; Table S2). Ecotype divergence accounted for for much of the variation of PC1 (*F*_*2,225*_ =211.04, *P* < 0.0001; Figure 2A; Table 1). Within ecotype, there was a significant effect of accession on PC1 (*F*_*8,225*_ = 23.70, *P* < 0.0001), and there was a significant effect of the GA treatment on PC1 (*F*_*1,225*_ = 15.22, *P* = 0.0001). While there was a significant accession x treatment effect for PC1 (*F*_*8,225*_ = 2.59, *P* = 0.0101), the treatment x ecotype effect was not significant (*P* = 0.1930). In contrast to PC1, both of the interactions were significant for PC2 (Eigenvalue = 5.632; accession x treatment: *F*_*8,225*_ = 7.61, *P* < 0.0001; ecotype x treatment: *F*_*2,225*_ = 6.16, *P* = 0.0025; Fig. 2A) and PC3 (Eigenvalue = 3.443; accession x treatment: *F*_*8,225*_ =4.03; *P* = 0.0002, ecotype x treatment: *F*_*2,225*_ = 42.07, *P* < 0.0001; Fig. 2B). Accession x treatment and ecotype x treatment interactions were not significant for PC4 (Table 1).

**Table 1.** Effects of individual factors and their interactions on the first four principal components of 37 morphological traits quantified in the experiment comparing ecotypes and accessions.

| Trait | Source | DF | SS | F | P |
| --- | --- | --- | --- | --- | --- |
| PC1 | Accession[Ecotype] | 8 | 730.32 | 23.70 | <0.0001* |
| PC1 | Ecotype | 2 | 1626.14 | 211.04 | <0.0001* |
| PC1 | Treatment | 1 | 58.63 | 15.22 | 0.0001* |
| PC1 | Accession*Treatment[Ecotype] | 8 | 79.71 | 2.59 | 0.0101* |
| PC1 | Ecotype*Treatment | 2 | 12.77 | 1.66 | 0.1930 |
| PC2 | Accession[Ecotype] | 8 | 366.72 | 20.82 | <0.0001* |
| PC2 | Ecotype | 2 | 54.38 | 12.35 | <0.0001* |
| PC2 | Treatment | 1 | 173.77 | 78.92 | <0.0001* |
| PC2 | Accession*Treatment[Ecotype] | 8 | 134.09 | 7.61 | <0.0001* |
| PC2 | Ecotype*Treatment | 2 | 27.12 | 6.16 | 0.0025* |
| PC3 | Accession[Ecotype] | 8 | 169.98 | 14.15 | <0.0001* |
| PC3 | Ecotype | 2 | 5.76 | 1.92 | 0.1492 |
| PC3 | Treatment | 1 | 51.18 | 34.08 | <0.0001* |
| PC3 | Accession*Treatment[Ecotype] | 8 | 48.37 | 4.03 | 0.0002* |
| PC3 | Ecotype*Treatment | 2 | 126.35 | 42.07 | <0.0001* |
| PC4 | Accession[Ecotype] | 8 | 143.39 | 16.28 | <0.0001* |
| PC4 | Ecotype | 2 | 109.37 | 49.66 | <0.0001* |
| PC4 | Treatment | 1 | 0.51 | 0.46 | 0.4981 |
| PC4 | Accession*Treatment[Ecotype] | 8 | 15.28 | 1.73 | 0.0916 |
| PC4 | Ecotype*Treatment | 2 | 0.42 | 0.19 | 0.8247 |

\*Significant after Bonferroni correction for multiple testing ( $P < 0.0125$ )

**Figure 2.**
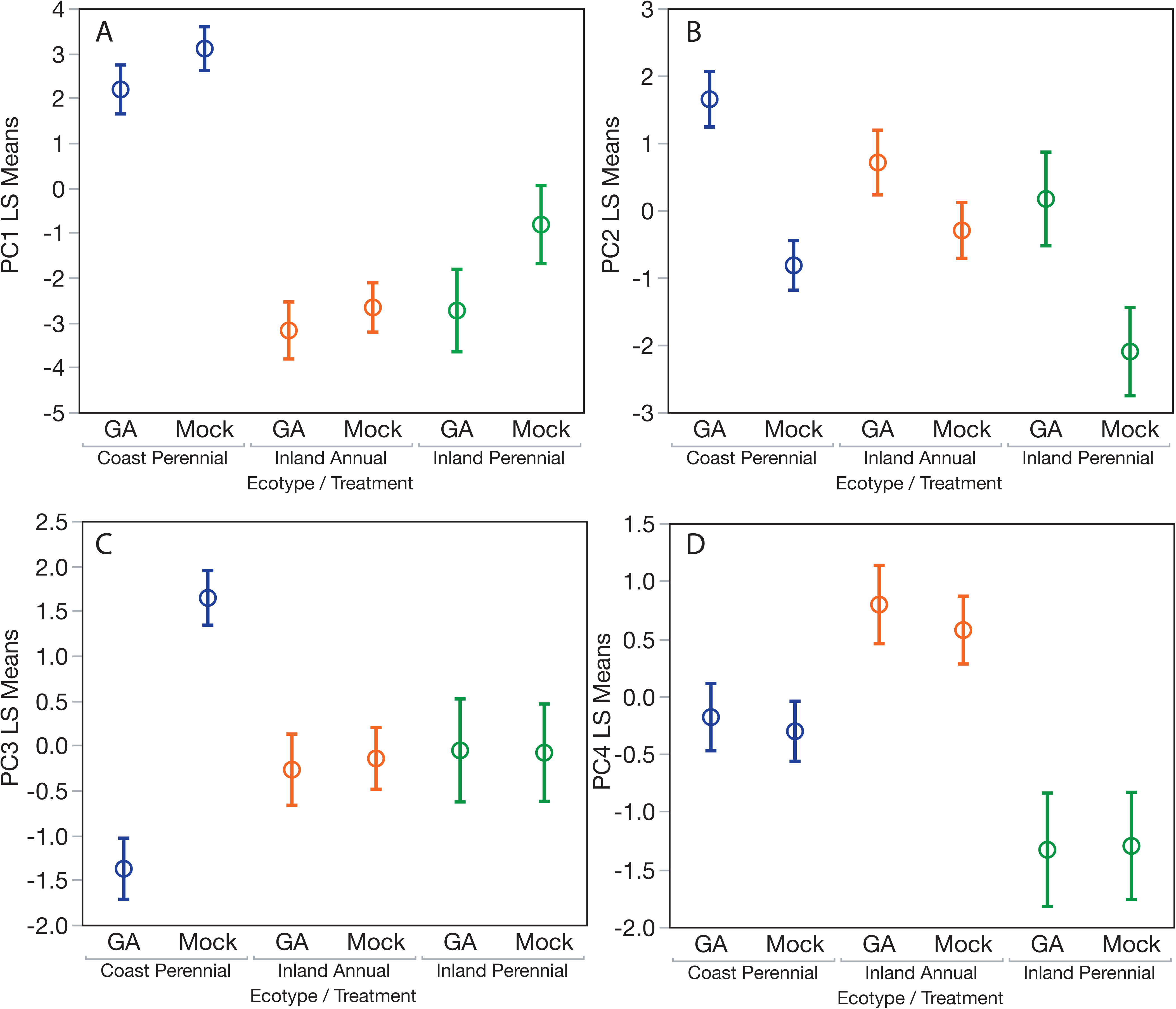
Least square means of ecotype x treatment interactions of the first four principal components for coastal perennial, inland annual, and inland perennial ecotypes in response to mock and gibberellin treatments. Error bars are standard errors.

Since there were significant ecotype x treatment interactions for PC2 and PC3, indicating a role of the GA pathway in ecotype divergence, we were interested in which traits might be driving these patterns. Second and third internode length, plant height, total number of stoloniferous nodes sprouting adventitious roots, total number of aerial branches, total number of stolons, and the length of the longest aerial branch all heavily loaded (> 0.40) onto PC2 (Table S2). The total number of aerial branches, the length of the longest aerial branch, number of nodes with adventitious roots, and leaf width heavily loaded onto PC3 (Table S2).

It should be noted that the effect of GA varied across accessions within ecotype. For example, the coastal PGR accession is the tallest coastal accession with the fewest stolons. This difference in overall morphology likely explains why its response to GA differed from the other four coastal accessions (Fig. S2). The inland perennials showed a similar level of response to GA as the coastal perennials for PC2, but not for PC3 (Fig. 2, S2).

### Interactions of GA application with the adaptive DIV1 inversion

Based on a Scree test (Fig. S3), we retained the first three PCs for analyses of the parental genotypes and NILs. Consistent with previous studies (Lowry and Willis 2010; Friedman 2014), the coastal perennial (SWB-S1) and inland annual (LMC-L1) lines were highly divergent in morphological traits and the two lines were differentiated strongly along PC1 (*F*_*1,163*_ = 314.39, *P* < 0.0001; Fig. 3A; Table 2). Similar to the accession analysis (above), the line x treatment interaction was not significant for PC1, but was highly significant for PC2 (*F*_*1,163*_ = 262.49, *P <* 0.0001; Table 2). The coastal perennial line (SWB-S1; *N* = 31 GA treatment, 42 Mock treatment) responded more strongly (Fig. 3B) to GA treatment for PC2 than the inland annual line (LMC-L1; *N* = 47 GA treatment, 47 Mock treatment), just as we found across coastal and inland populations more generally (above). The set of traits heavily loading (>0.40) onto PC2 was very similar to that for PC2 of the accession analysis (above): Third internode length, plant height, total number of stoloniferous nodes sprouting adventitious roots, number of adventitious roots at the first node, total number of aerial branches, total number of stolons, and the length of the longest aerial branch (Table S3).

**Table 2.** Effects of line (SWB-S1 vs. LMC-L1), treatment (GA vs. Mock), and the interaction on the first three principal of 37 morphological traits quantified for the parental inbred lines.

| Trait | Source | SS | F | P |
| --- | --- | --- | --- | --- |
| PC1 | Line | 2027.14 | 314.39 | <0.0001* |
| PC1 | Treatment | 145.75 | 22.6 | <0.0001* |
| PC1 | Line*Treatment | 7.06 | 1.10 | 0.2969 |
| PC2 | Line | 42.25 | 15.61 | 0.0001* |
| PC2 | Treatment | 257.86 | 95.24 | <0.0001* |
| PC2 | Line*Treatment | 262.49 | 96.95 | <0.0001* |
| PC3 | Line | 2.12 | 0.63 | 0.4271 |
| PC3 | Treatment | 32.51 | 9.74 | 0.0021* |
| PC3 | Line*Treatment | 15.73 | 4.71 | 0.0314 |

\*Significant after Bonferroni correction for multiple testing ( $P < 0.0083$ )

**Figure 3.**
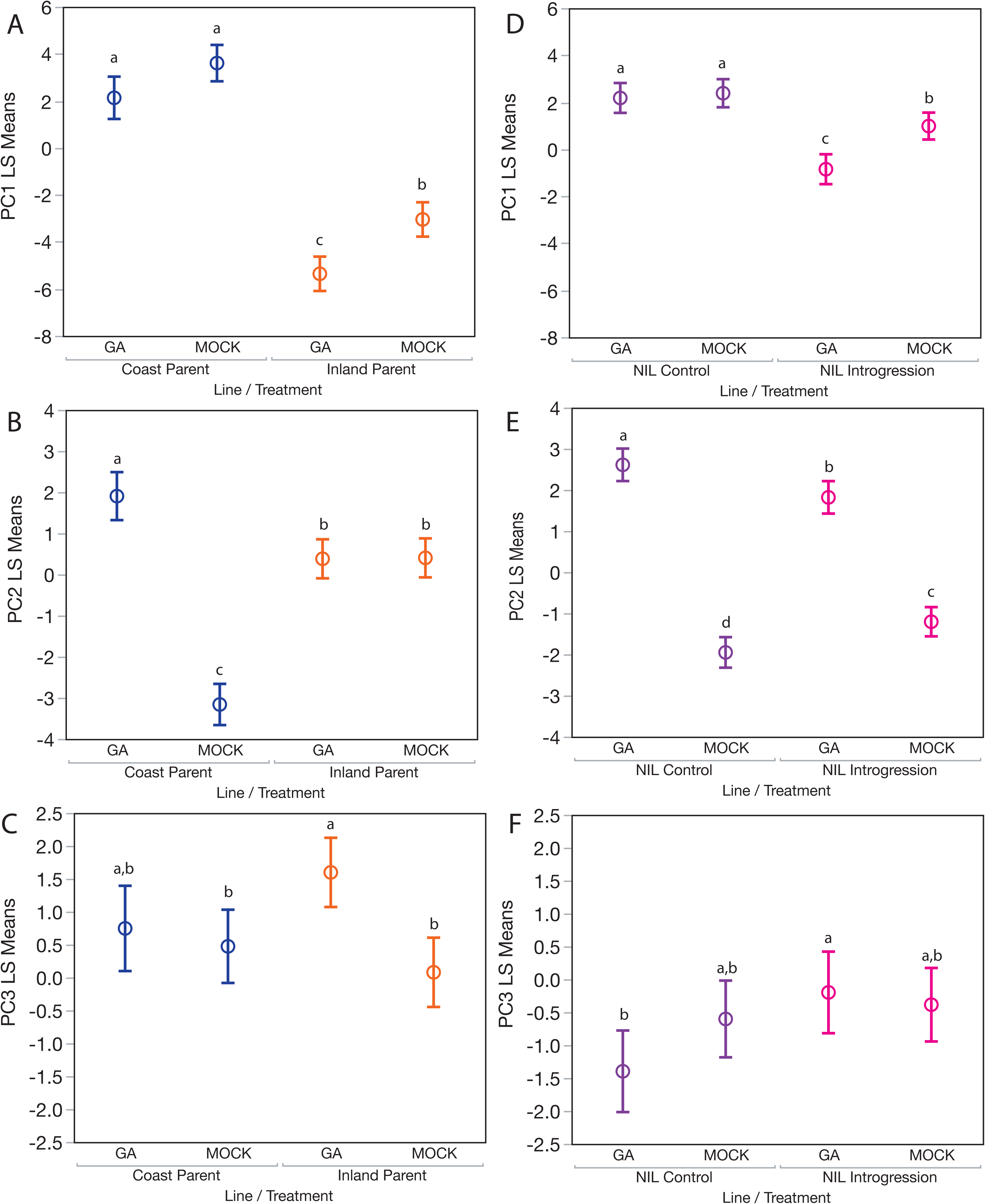
Least square means of line x treatment interactions of coastal perennial (SWB-S1) and inland annual (LMC-L1) parents and coastal perennial genetic background near-isogenic lines (NILs) to mock and gibberellin treatments. The introgression NILs were homozygous for the inland annual orientation of the *DIV1* inversion, while the control NILs were homozygous for the coastal perennial orientation of the *DIV1* inversion. Lower case letters indicate significant differences of least square means by Tukey HSD *post hoc* analyses. Error bars are standard errors.

The *DIV1* chromosomal inversion is one of many loci responsible for divergence between the annual and perennial ecotypes (Hall et al. 2006; Lowry & Willis 2010; Friedman 2014). Thus, main effects and interactions in the NILs were expected to be subtler than for the parental lines. As in previous studies (Lowry and Willis 2010; Friedman 2014), the inversion had major effects on traits associated with ecotype divergence, with highly significant main effects on PC1 (*F*_*1,166*_= 51.23, *P <* 0.0001; Table 3). The GA treatment had significant effects on PC1 (*F*_*1,166*_ = 10.86, *P* = 0.0012) and PC2 (*F*_*1,166*_ = 606.48, *P <* 0.0001). There was a significant, line x treatment interactions for PC2 (*F*_*1,166*_ = 24.91, *P* < 0.0001; Fig. 3E), where Introgression-NILs (*N* = 39 GA treatment, 48 Mock treatment) containing perennial inversion orientation responded more to the GA treatment than the Control-NILs (*N* = 39 GA treatment, 44 Mock treatment) containing the annual orientation. None of the effects were significant for PC3 after Bonferroni correction for multiple testing (Table 3).

**Table 3.** Effects of line (Introgression-NIL vs. Control-NIL), treatment (GA vs. Mock), and the interaction on the first three principal of 37 morphological traits quantified for the parental inbred lines.

| Trait | Source | SS | F | P |
| --- | --- | --- | --- | --- |
| PC1 | Line | 206.86 | 51.23 | <0.0001* |
| PC1 | Treatment | 43.84 | 10.86 | 0.0012* |
| PC1 | Line*Treatment | 28.17 | 6.98 | 0.0090 |
| PC2 | Line | 0.02 | 0.01 | 0.9072 |
| PC2 | Treatment | 606.48 | 389.98 | <0.0001* |
| PC2 | Line*Treatment | 24.91 | 16.02 | <0.0001* |
| PC3 | Line | 21.12 | 5.50 | 0.0202 |
| PC3 | Treatment | 3.90 | 1.02 | 0.315 |
| PC3 | Line*Treatment | 10.15 | 2.64 | 0.1059 |

\*Significant after Bonferroni correction for multiple testing ( $P < 0.0083$ )

### Effects of the DIV1 inversion on resistance compound concentrations

We quantified the concentrations of seven PPGs (Table 4). Consistent with our previous observations (Holeski et al. 2013), the coastal perennial parental (SWB) plants produced 2.5 times more total PPGs than the inland annual parental (LMC) plants (*F*_*1,31*_ = 51.03; *P* < 0.0001; Table 4; Fig. 4). There were also significant differences for six out of seven of the PPGs between the coastal perennial (SWB) and inland annual (LMC) parental lines.

**Table 4.** Means (standard errors) and Tukey HSD post-hoc results for the concentrations of phenylpropanoid glycosides in coastal perennial (SWB) and inland annual (LMC) parents as well as near-isogenic lines of the *DIV1* inversion in the coastal perennial genetic background.

| Trait | Coastal Parent (N=25) | Inland Parent (N=8) | Control NIL (N=54) | Introgression NIL (N=35) |
| --- | --- | --- | --- | --- |
| Total PPGs | 164.24 (7.74)A | 65.65 (13.69)C | 161.32 (5.27)A | 119.67 (6.55)B |
| Conandroside | 105.11 (6.65)A | 42.26 (11.75)B | 106.98 (4.52)A | 55.71 (5.62)B |
| Calceolarioside A | 37.15 (2.13)B | 12.79 (3.77)C | 32.77 (1.45)B | 46.30 (1.80)A |
| Mimuloside | 7.93 (0.64)A | 3.62 (1.13)B | 7.54 (0.43)A | 4.24 (0.55)B |
| Verbascoside | 6.91 (0.36)A | 1.90 (0.63)B | 6.89 (0.24)A | 5.90 (0.30)A |
| Calceolarioside B | 0.98 (0.08)A | 0.33 (0.13)B | 0.66 (0.06)B | 0.62 (0.06)B |
| Unknown PPG 16 | 5.52 (0.44)A | 4.56 (0.77)A | 5.87 (0.30)A | 6.09 (0.37)A |
| Unknown PPG10 | 0.64 (0.04)B | 0.19 (0.07)C | 0.60 (0.03)B | 0.81 (0.03)A |

Mean (SE) and Tukey HSD post-hoc results. Sample sizes in parentheses.

**Figure 4.**
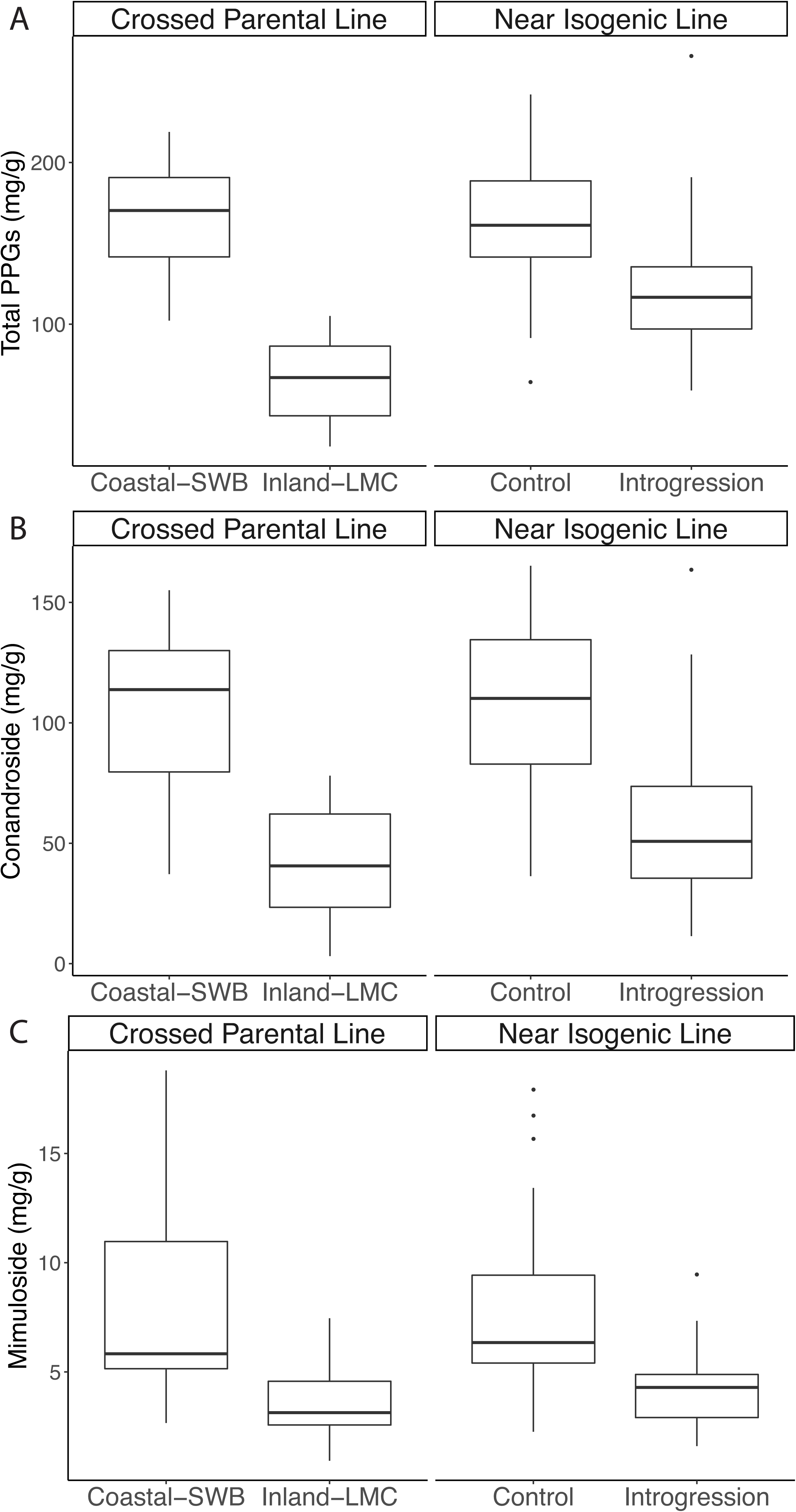
Concentration of phenylpropanoid glycosides for coastal perennial (SWB-S2xS3) and inland annual (LMC-L2xL3) parents as well as coastal perennial genetic background near-isogenic lines (NILs). The introgression NILs were homozygous for the inland annual orientation of the *DIV1* inversion, while the control NILs were homozygous for the coastal perennial orientation of the *DIV1* inversion.

Analysis of the *DIV1* NILs revealed that the introgressed region containing the inversion had major effects on foliar concentrations of PPGs. Control NILs that were homozygous for the coastal orientation of the *DIV1* inversion produced 35% higher concentrations of total PPGs than the introgression NILs, which were homozygous for the inland *DIV1* orientation (*F*_*1,87*_ = 22.70; *P* < 0.0001). In addition, the *DIV1* locus had significant effects on four out of the seven individual PPGs. Interestingly, the control NILs had higher concentrations of conandroside and mimuloside, but lower concentrations of calceolarioside A and unknown PPG10, than the introgression NILs (Table 4). Thus, the *DIV1* inversion influences both the total concentration of PPGs as well as the composition of suites of these PPGs.

## DISCUSSION

In this study, we identified a potential mechanism underlying a coordinated evolutionary shift between growth, reproduction, and herbivore resistance in the *M. guttatus* species complex. Our finding that perennial accessions were more responsive to the GA treatment along the second principal component axis (PC2; Fig. 2B) is consistent with the hypothesis that perennial populations have less GA biosynthesis and/or activity. This pattern was replicated for the coastal perennial and inland annual parents of the NILs (Fig. 3B). Further, the finding that the introgression NILs had less response to GA addition along the PC2 axis than the control NILs (Fig. 3E) is consistent with the hypothesis that the perennial orientation of the *DIV1* inversion reduces GA biosynthesis/activity. The morphological traits most likely impacted through this mechanism include plant height, adventitious root production, and whether lateral branches develop as vegetative stolons or upcurved reproductive shoots. Finally, we found that coastal perennial plants and NILs containing the perennial orientation of the inversion produced higher concentration of defensive PPG compound than inland annual plants and NILs containing the inland orientation of the inversion. Therefore, the coastal perennial orientation of the *DIV1* inversion promotes allocation to long-term growth and herbivore resistance, while the inland orientation promotes allocation of resources to traits associated with rapid reproduction. We discuss these findings in the context of the broader literature below.

### Environmental gradients and the evolution of growth, reproduction, and defense traits

Studies of intraspecific variation among natural populations adapted to different soil water availability regimes provide an excellent opportunity to understand how the abiotic environment influences the relative allocation of resources by plants to growth and constitutive/induced resistance. Soil water is one of the most limiting factors for plants on Earth (Whittaker 1975; Bohnert et al. 1995; Bray 1997) and can drastically differ in availability among seasons (Cowling et al. 1996), which in turn influences plant resource allocation (Juenger 2013). The timing of soil water availability can dictate the length of the growing season. One major evolutionary strategy for seasonally low water availability is to allocate resources primarily to growth and reproduction to achieve an early flowering, drought escape life-history (Ludlow 1989; Juenger 2013; Kooyers 2015). Beyond selection on plants, soil moisture gradients can drive the abundance of herbivores, which in turn exert their own selective pressures (Kooyers et al. 2017).

In *M. gutattus*, evolutionary shifts across a soil moisture gradient drives changes in the allocation not only between growth and reproduction, but also for herbivore resistance (Lowry et al. 2008; Holeski et al. 2013; Kooyers et al. 2017). The phenotypic differences between coastal perennial and inland annual populations is likely driven by multiple selective pressures that are tied to the divergence in soil water availability between coastal and inland habitats. Inland annual habitats generally dry out very quickly at the end of the spring, which leaves little time for a plant to reproduce before being killed by the summer drought. Further, the short growing season may also prevent the establishment of sizable herbivore populations, which would explain the low level of leaf damage in fast drying inland annual habitats (Lowry et al. 2008; Kooyers et al. 2017). In contrast, the year-round soil water availability of coastal habitats means that plants growing there have much more time to allocate resources to vegetative growth and herbivore resistance. In addition, wet coastal habitats can build up a considerable load of herbivores, which is likely contribute to the higher incidences of leaf damage and early season mortality at the coast (Lowry et al. 2008; Lowry and Willis 2010; Popovic & Lowry 2019). For intraspecific differences among populations, the strength of herbivore pressure is thought to be a key driver of plant resistance (Hahn and Maron 2016). It should be noted that a fair amount of leaf damage in coastal habitats may also be due to oceanic salt spray (Boyce 1954; Ahmed and Wainwright 1976; Griffiths 2006; Lowry et al. 2009; Popovic & Lowry 2019). However, a recent manipulative field experiment did find much greater levels of herbivory of *M. guttatus* plants in common gardens in coastal habitat than inland habitat (Popovic & Lowry 2019).

Our findings in *M. guttatus* are likely to have implications for intraspecific variation in many other plant species as well. There are many studies that have found similar developmental differences between coastal and inland populations as we have found for *M. guttatus* (reviewed in Lowry 2012). Given the commonality of coastal plants investing more heavily in lateral vegetative branches versus inland populations investing primarily in upright flowering branches, we predict that coastal population of plants will generally be more highly defended than inland populations, particularly in Mediterranean climates with steep soil water availability gradients.

### The role of pleiotropy and linkage

The results of this study and previous studies (Lowry and Willis 2010; Friedman 2014; Friedman et al. 2015) collectively demonstrate that adaptive chromosomal inversion *DIV1* contributes to the shift in allocation between long-term growth, short-term fecundity, and herbivore resistance. An outstanding question is whether this coordinated shift is due to genetic linkage or pleiotropy. Chromosomal inversions are thought to evolve as adaptation “supergenes,” which can trap multiple linked adaptive loci through their suppression of meiotic recombination (Dobzhansky 1970; Kirkpatrick and Barton 2006; Schwander et al. 2014; Wellenreuther and Bernatchez 2018). Thus, the fact that the *DIV1* inversion contributes to the evolution of multiple phenotypes could be result of adaptive alleles at multiple linked loci being held together in tight linkage by the chromosomal inversion. Alternatively, a single gene within the inversion could have pleiotropic effects on all of the phenotypic changes. Functional studies involving transformation or gene editing of candidate genes within the inversion will be necessary to distinguish between these alternative hypotheses.

Two other recent studies have also found potential pleiotropic effects of genes on allocation to reproduction and herbivore resistance. Rasmann et al. (2018) found that NILs containing genetic variants of the *Flowering Locus C* (*FLC*) gene in *Cardamine hirsute* are responsible for a trade-off between early flowering and herbivore resistance in terms of glucosinolate production. Kerwin et al. (2015) found that there was a positive correlation in *Arabidopsis thaliana* between glucosinalate production and flowering time for mutant alleles of genes in the glucosinolate biosynthetic pathway. Overall, both of these studies identified the same trade-off of rapid reproduction versus herbivore resistance that we found in our study, although mediated through independent genetic mechanisms.

### A hormonal basis of a coordinated shift in the evolution of growth, reproduction, and herbivore resistance?

The finding that coastal perennial and inland annual plants respond differentially to GA is consistent with the role of this hormone playing a role in their evolutionary divergence. A recent outlier analysis of coastal perennial and inland annual populations of *M. gutattus* found that the gene *GA20-oxidase2* (*GA20ox2*) was a major allele frequency outlier between the ecotypes within the vicinity of the *DIV1* inversion (Gould et al. 2017). This gene is a strong potential candidate gene for a pleiotropic shift in allocation between growth and reproduction. GA-oxidases are involved in the evolution of dwarfed coastal populations of *A. thaliana* (Barboza *et al*. 2013) and played a key role in the development of dwarfed Green Revolution rice and barley (Sasaki et al. 2002; Jia et al. 2009). Further, the DELLA gene, *GAI*, is located within the vicinity of another growth regulatory QTL (*DIV2*). *GAI* was also an allele frequency outlier between coastal and inland populations (Gould et al. 2017). Friedman (2014) found that the *DIV1* and *DIV2* loci interact epistatically. Thus, it would not be surprising if the genetic changes that underlie both QTL are in the same molecular pathway. Further, the negative antagonism between the GA and JA hormone pathways via the DELLA-JAZ signaling node (Havko et al. 2016; Guo et al. 2018; Howe et al. 2018) suggests a direct mechanism by which trade-offs between growth, reproduction, and resistance could easily evolve. Future functional genetic studies will be needed to determine whether these genes in fact are involved in adaptive shifts between growth, reproduction, and defense underlying local adaptation in this system.

While we saw a greater response to GA addition in coastal plants and observed an interaction between GA addition and the inversion, other hormones could also play a role or even be the ultimate cause of the divergence between the coastal perennial and inland annual ecotypes. Three major classes of hormones, Auxins, Brassinosteroids and Gibberellins, are all associated with growth phenotypes like those that differ between coastal perennial and inland annual ecotypes of *M. guttatus* (Ross and Quittenden 2016; Unterholzner et al. 2016). These hormones interact in multiple ways, which have yet to be fully elucidated, to result in shifts in growth/reproduction phenotypes.

### Conclusions and future directions

There are numerous evolutionary and ecological models that make predictions on the evolution of relationships between growth, reproduction, and herbivore resistance. While recent meta-analyses have found that some models have moderate support (Endara and Coley 2011), there are many exceptions and some models appear to not be well supported at all (Stamps et al. 2003; Hahn and Maron 2016; Smilanich et al. 2016). The reasons that these models do not hold up are often attributed to vast variation in the extrinsic environmental factors that exert selective pressures on plant populations, broad variation in life-history among plant species, differences between interspecific and intraspecific variation, and variation in herbivore abundances across environmental gradients (Stamps et al. 2003; Hahn and Maron 2016; Smilanich et al. 2016; Hahn et al. 2019). Less well appreciated are the molecular genetic mechanisms that underlie shifts in allocation between growth, reproduction, and herbivore resistance (Kerwin et al. 2015; Rasmann et al. 2018). The nature of the gene networks responsible for shifts in allocation may also be very important for whether or not particular systems will conform to a given evolutionary or ecological model. Future research should focus on uncovering the molecular mechanisms that underlie the evolution of growth, reproduction, and defense trade-offs in natural populations and integrate predictions from those mechanisms into ecological and evolutionary models.

## Supplemental Materials

**Table S1.** Collection locations of population accessions used in this study, including the number of individuals used for ANOVAs of principal components per each accession.

**Table S2.** Trait loadings on the first four principle components axes for experiment comparing population accession. Percent of variation explained by each PC included in parentheses.

**Table S3.** Trait loadings on the first three principle components axes for all plants grown in the experiment with near isogenic lines and the parental genotypes. Percent of variation explained by each PC included in parentheses.

---

**Figure S1.**
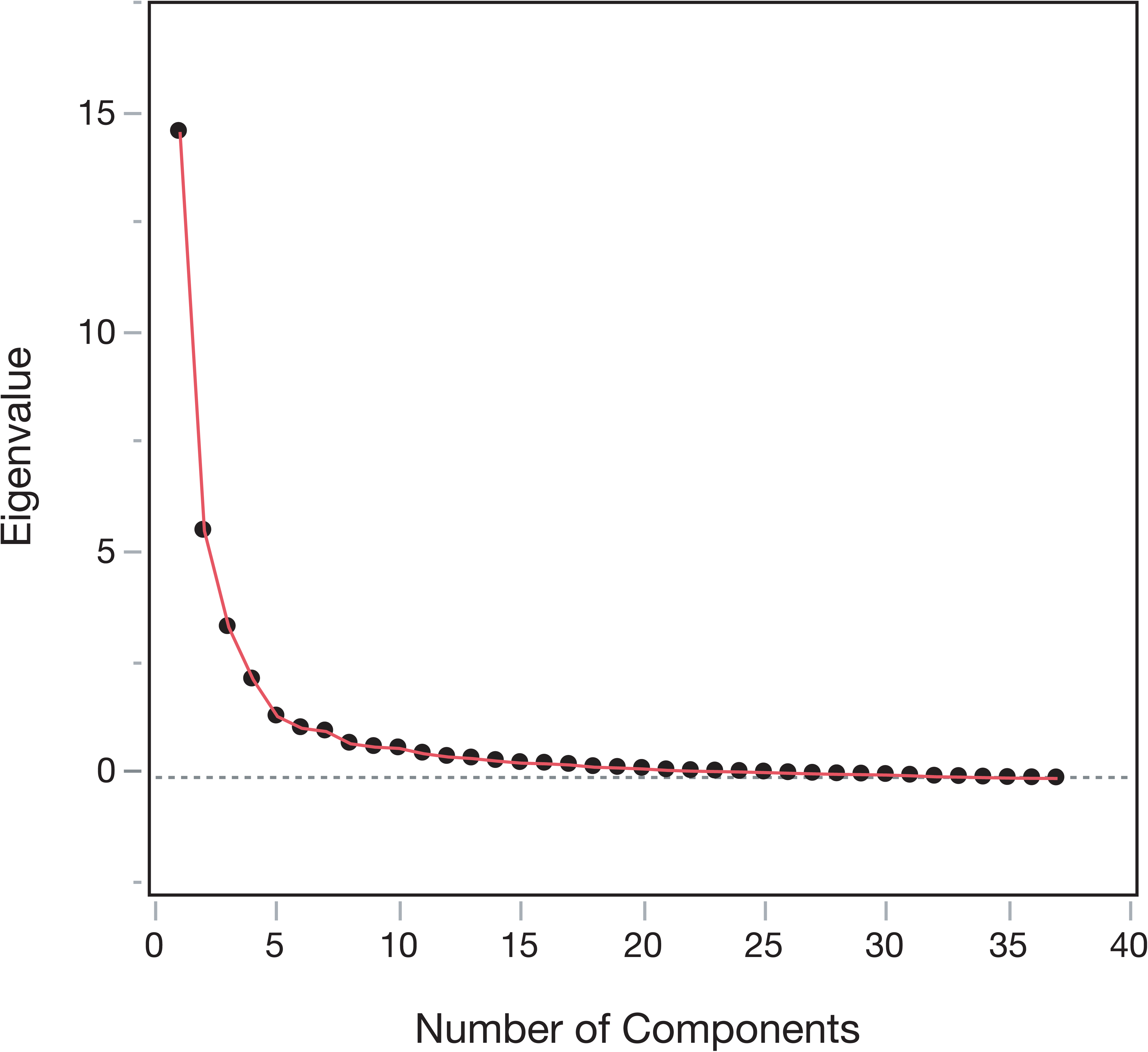
Scree plot of eigenvalues for principal components analysis of trait data collected for population accessions

**Figure S2.**
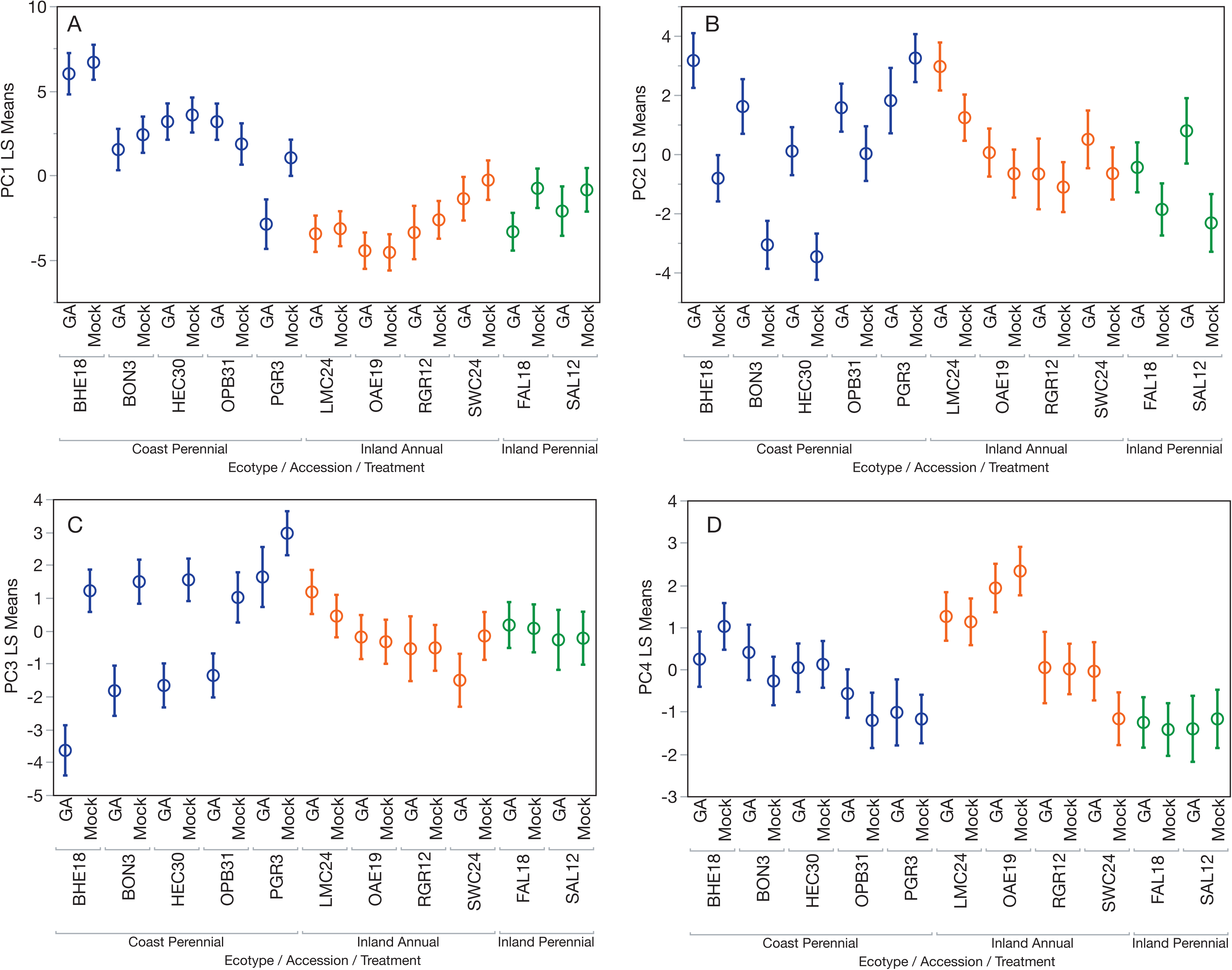
Least square means of accession x treatment interactions of the first four principal components for coastal perennial, inland annual, and inland perennial accessions in response to mock and gibberellin treatments.

**Figure S3.**
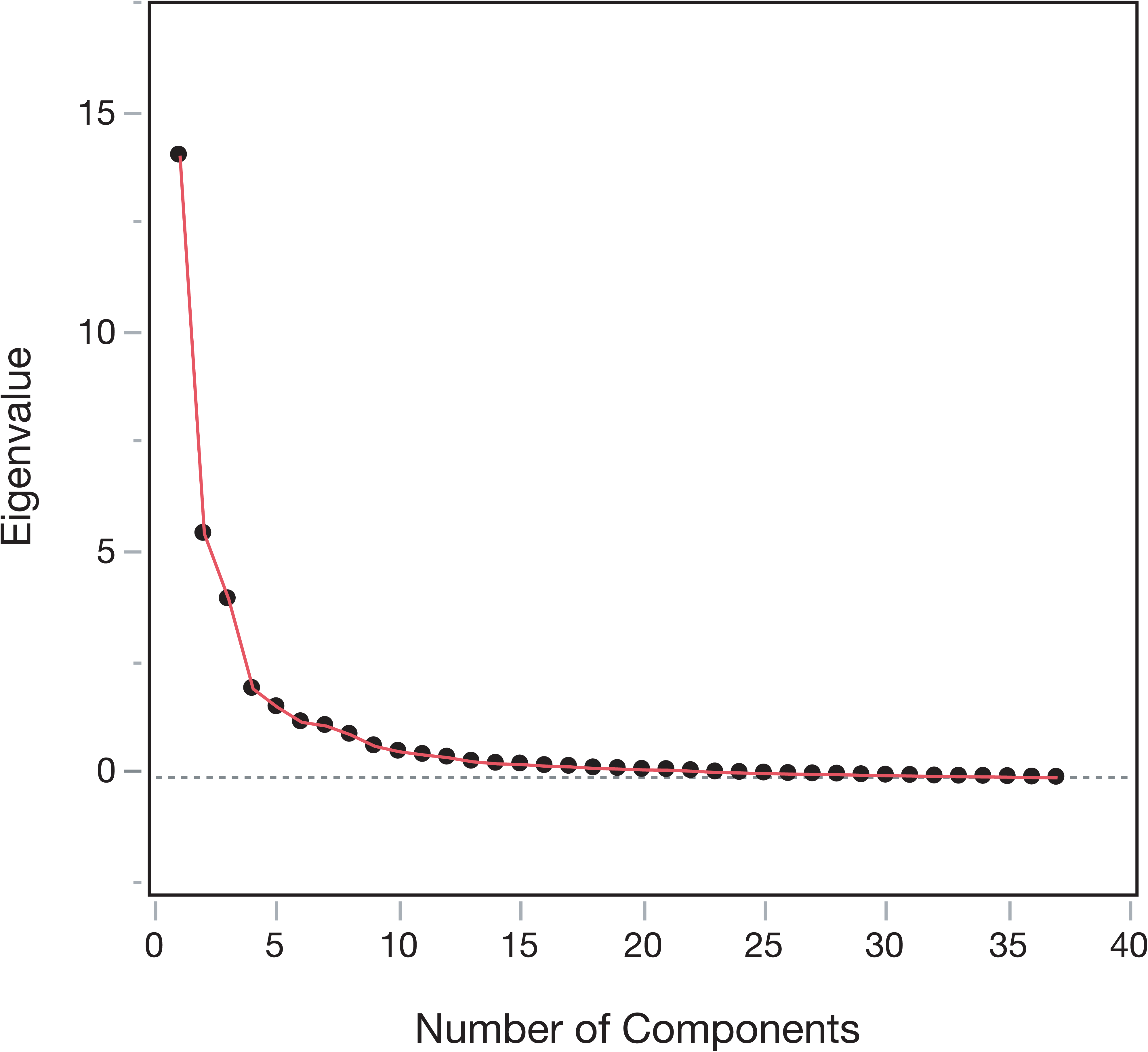
Scree plot of eigenvalues for principal components analysis of trait data collected for experiment with near isogenic lines and parental lines.

Author Contributions
DL and DP designed the experiments; DL, DP and DB conducted the experiments; LH conducted the analyses of herbivore resistance compounds; DL and LH wrote the manuscript

## Acknowledgements

We would like to thank Sol Chavez for assisting with the quantification of PPGs. Seed collections were originally made possible by permission from the state parks of Oregon and California. Funding for this research was provided by Michigan State University through a startup package to DBL.

Raw data from experiments will be deposited at Dryad upon publication

